# Very high-density platelets determine reactivity and activity of circulating platelets

**DOI:** 10.1101/388744

**Authors:** P. Järemo

## Abstract

**Background:** For many decades, platelets have been known to display a substantial density heterogeneity. Knowledge about the origins and functions of specific platelet density subpopulations is scarce. This study investigates if very high-density (1.09 kg/L) platelets impact upon the reactivity and activity of all platelets.

**Methods:** Subjects (*n*=52) were recruited in conjunction with a nurse-guided blood pressure control. Platelet reactivity in citrate anticoagulated whole blood, i.e. surface-bound P-selectin after provocation, was analysed using a flow cytometry technique. ADP (8.5 μmol/L) was employed as an agonist. Subsequently, the entire population was separated according to density into 17 subpopulations, with fraction 1 containing the densest platelets (1.09 kg/L). In each subfraction surface P-selectin expression was determined. Subjects were then divided according to the number of very high-density platelets in 1.09^high^ (*n*=17) and 1.09^low^ (*n*=35) demonstrating >8×10^9^/L and ≤8×10^9^/L 1.09 kg/L fraction 1 platelets, respectively.

**Results:** Surface-attached P-selectin after provocation reveals that 1.09^high^ associates with increased whole blood reactivity of the entire platelet population. The level of significance was *p*≤0.01 (8.5 μmol/L ADP). Furthermore, 1.09^high^ relates to increased spontaneous activity of density populations, as evidenced by membrane-bound P-selectin. For the fractions *nos*. 2, 4-7, 9, 10 the differences were significant, with *p*-values ranging from *p*≤0.05 to *p*≤0.01.

**Conclusion:** The number of very high-density (1.09 kg/L) platelets reflects the reactivity of the entire population. It is also closely related to subfraction P-selectin activity. It is unlikely that platelets gain density when circulating. Therefore, evidence suggests that very dense cells are created for this purpose at thrombopoiesis. It is tenable that such platelets regulate the reactivity of the entire population.

## Introduction

Circulating platelets exhibit a substantial heterogeneity with respect to density. The differences are believed to originate at the fragmentation of megakaryocytes in the bone marrow [1]. High-density platelets contain more cell organelles and glycogen [2], [3], display enhanced metabolism [4] and adhere more rapidly [5]. When activating, platelets undergo morphological changes and turn on the major glycoprotein IIb/IIIa (GP IIb/IIIa) receptor, which binds fibrinogen. Furthermore, their α- and dense granules then fuse with the cell membrane, thereby releasing a cargo of platelet-specific proteins such as P-selectin and β-thromboglobulin [6]. Platelets have a multitude of functions and frequently act in cooperation with other cells [7], [8]. Examples of this include creating a barrier that limits blood loss at injured vasculature, providing a surface for thrombin generation, orchestrating inflammatory reactions through surface P-selectin receptors and endowing wound healing by rebuilding damaged tissue [8]. It is unlikely that every platelet takes on all these assignments.

During the last few decades, most studies have assumed that platelets are homogeneous, but interest in platelet heterogeneity has risen with the discovery of coated platelets [8], [9], [10]. These appear in the test tube in response to dual actions of collagen and thrombin. Procoagulant platelets are smaller, display more surface-bound P-selectin but less fibrinogen than their aggregating counterparts. In this laboratory, heterogeneity of resting platelets has been extensively investigated [11] and no subpopulation display procoagulant features. As judged from surface-attached fibrinogen, very high-density populations circulate more activated than neighbouring less dense cells [12], peak density platelets are activated to a lesser extent, whereas lighter platelets again display more activity markers [13].

Platelet density alterations are features of disease state. Examples include that density increases in conjunction with acute myocardial infarctions [14] and associates inversely with inflammatory reactions [15]. Small high-density platelets are features of active inflammatory bowel disease [16] and higher peak platelet density is a characteristic of diabetes type 2 [17]. In contrast, low platelet density characterises essential thrombocythemia [18] [19] and preeclampsia [20]. Changes of the activation pattern of platelet density subpopulations are features of Alzheimer’s disease and then the bulk of platelets show less surface-bound fibrinogen [21]. Platelet heterogeneity is also important in malignancy in that a tiny proportion of larger platelets predicts prostatic cancer recurrence [22].

Platelet density diversity has been investigated for many years, but information about the functions of platelet density fractions is incomplete. The current study thus examines if very high-density cells (1.09 kg/L) affect reactivity of the entire platelet population.

## Methods

Subjects (*n*=52) were recruited as they attended a nurse-guided blood pressure control (Table 1). Automated hemograms were determined electronically and glycosylated haemoglobin (HbA1c) by a standard procedure. Citrate anticoagulated whole blood was used for analysing reactivity and activity of all platelets and, to minimise deviations, due to sample aging the determinations were commenced approximately 2 hours after sampling. The number (%) of cells expressing P-selectin/ fibrinogen was visualised using a Beckman Coulter EPICS XL-MCL™ flow cytometer (Beckman Coulter, Inc., Brea, Cal, USA) [23], [24]. Platelets were identified with an antibody towards glycoprotein Ib. A non-stimulated control containing EDTA was subtracted from the experimental values. The following antibodies were used.

**Table 1.**
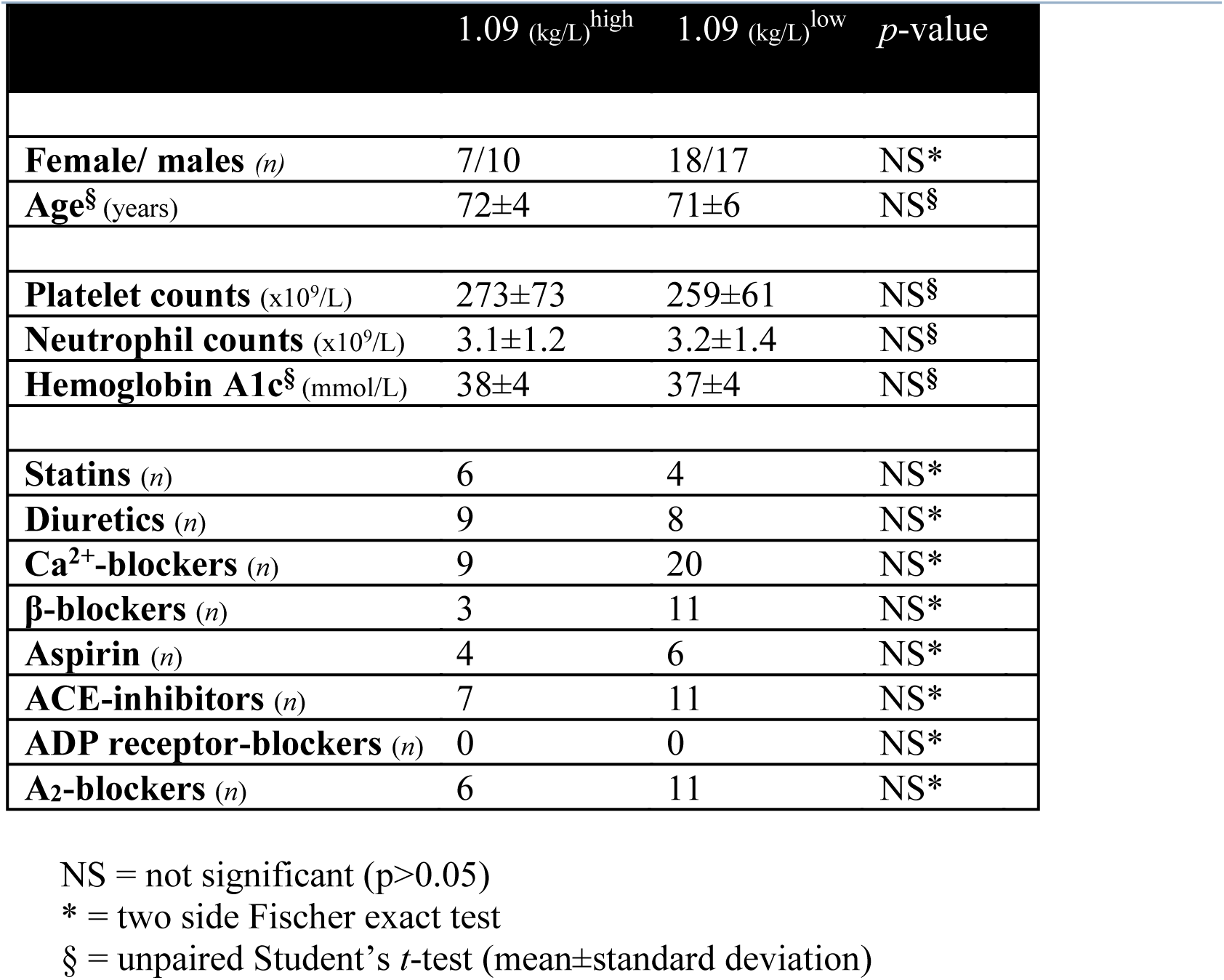
Baseline characteristics for individuals having higher (*n*=17) and lower (*n*=35) counts of very high-density (1.09 kg/L) platelets, respectively.

**Fibrinogen:** a polyclonal antihuman chicken antibody (Diapensia, Linköping, Sweden).

**Glycoprotein Ib**: a mouse monoclonal antibody (Dako AS, Glostrup, Denmark).

**P-selectin:** a monoclonal IgG1 mouse antibody (BD Biosciences, Sparks, MD, USA).

Platelet reactivity, i.e. their P-selectin and fibrinogen responses, was analysed using ADP (1.7 and 8.5 μmol/L) (Sigma-Aldrich, St Louis, Mo, USA) as agonists [24]. Subsequently, platelets were separated according to density utilising linear Percoll™ gradients covering the density span 1.09 - 1,04 kg/L manufactured essentially as previously described [11], [12], [21]. To avoid *in vitro* activity an inhibitory cocktail containing theophylline, prostaglandin E^1^ and EDTA was employed as anticoagulant [11]. The platelet population was subsequently separated into density fractions (*n*=17) [11], [21]. In all fractions determination of platelet counts was carried out together with platelet P-selectin and fibrinogen membrane expressions. Finally, participants were divided according to the number of very high-density (1.09 kg/L) platelets. Group 1.09^high^ (*n*=17) showed more of such platelets (>8×10^9^/L) whereas the 1.09^low^ group had fewer (≤8×10^9^/L) (*n*=35). The two-side Fischer exact test and the unpaired Student’s *t*-test were used as statistics. *P-*values ≤ 0.05 were regarded as significant. Ethical permission was obtained from the Local Institution Review Board.

## Results

The quantity of 1.09 kg/L platelets was unaffected by demographic features and routine laboratory (i.e. platelet and neutrophil counts together with HbA1c) (Table 1). After provocation, 1.09^high^ individuals displayed significantly higher whole blood platelet reactivity, as estimated from surface-bound P-selectin (Table 2). The levels of significance when using ADP (1.7 and 8.5 μmol/L) as platelet agonists were *p*≤0.05 and *p*≤0.01, respectively. In contrast, the number of 1.09 kg/L platelets did not associate with surface-attached fibrinogen after stimulation as a blood reactivity measure. Furthermore, unprovoked membrane adjoined P-selectin failed to show differences between groups, but 1.09^high^ displayed reduced platelet activity *in vivo* as judged from fibrinogen binding in whole blood (*p*≤0.05) (Table 2).

**Table 2.** Platelet reactivity and activity (mean±SD) in whole blood of individuals having higher (*n*=17) and lower (*n*=35) concentrations of very high-density (1.09 kg/L) platelets, respectively.

|  | 1.09 kg/L<sup>high</sup> | 1.09 kg/L<sup>low</sup> | *p*-value |
| --- | --- | --- | --- |
| **Platelet-bound P-selectin after agonist provocation** |  |  |  |
| ADP (1.7 µmol/L) (%) | 17±9 | 12±8 | ≤0.05 |
| ADP (8.5 µmol/L) (%) | 46±16 | 35±16 | ≤0.01 |
| **Platelet-bound fibrinogen after agonist provocation** |  |  |  |
| ADP (1.7 µmol/L) (%) | 11±6 | 13±7 | NS |
| ADP (8.5 µmol/L) (%) | 42±16 | 40±16 | NS |
| **Platelet-bound P-selectin** |  |  |  |
| without agonist (%) | 7±4 | 6±3 | NS |
| **Platelet-bound fibrinogen** |  |  |  |
| without agonist (%) | 4±1 | 5±1 | ≤0.05 |

NS = not significant (Student *t*-test ( $p>0.05$ ))
% = percentage positive cells (either P-selectin or fibrinogen)

Figure 1 shows the platelet distribution in the gradients. For the 1.09^high^ group cell density was shifted to the right i.e. the mean density peak was decreased. Consequently, they demonstrated lower platelet counts in density fractions *nos.* 7, 8 (*p*≤0.05) and more platelets in fractions *nos*. 10, 11 (p≤0.05) (Figure 1). In addition, subpopulation P-selectin activity of 1.09^high^ group was enhanced (Figure 2). The *p*-values ranged from *p*≤0.05 (fractions *nos*. 2, 4, 5, 9, 10) to *p*≤0.01 (fractions *nos*. 6, 7). In contrast, spontaneous platelet-bound fibrinogen of lighter cells behaved differently in that 1.09^high^ subjects demonstrated less of the activity marker (Figure 3). The latter dissimilarities were significant for fractions *nos.* 11, 12 (*p*≤0.05).

**Figure 1.**
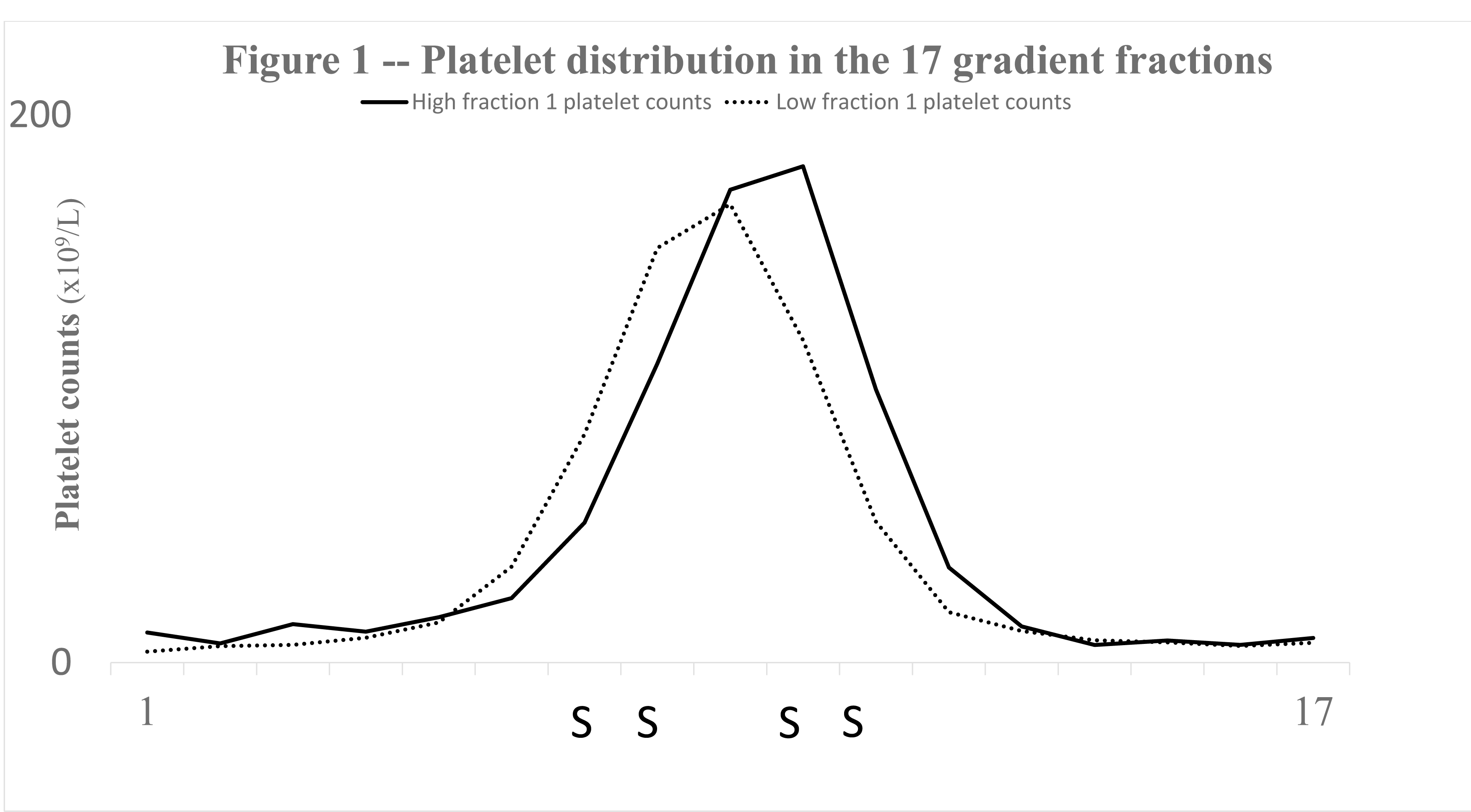
Shows, for the two groups (1.09^high^ and 1.09^low^), the distribution of platelets (x10^9^/L) (vertical axis) of the 17 density fractions of the gradients (horizontal axis). Significant differences are denoted with the character S. The platelet distribution of 1.09^high^ individuals was shifted to the right and subpopulations *nos.* 7, 8 contained fewer platelets (*p*≤.0.05) whereas the platelet counts of fractions *nos.* 10, 11 were increased (*p*≤.0.05).

**Figure 2.**
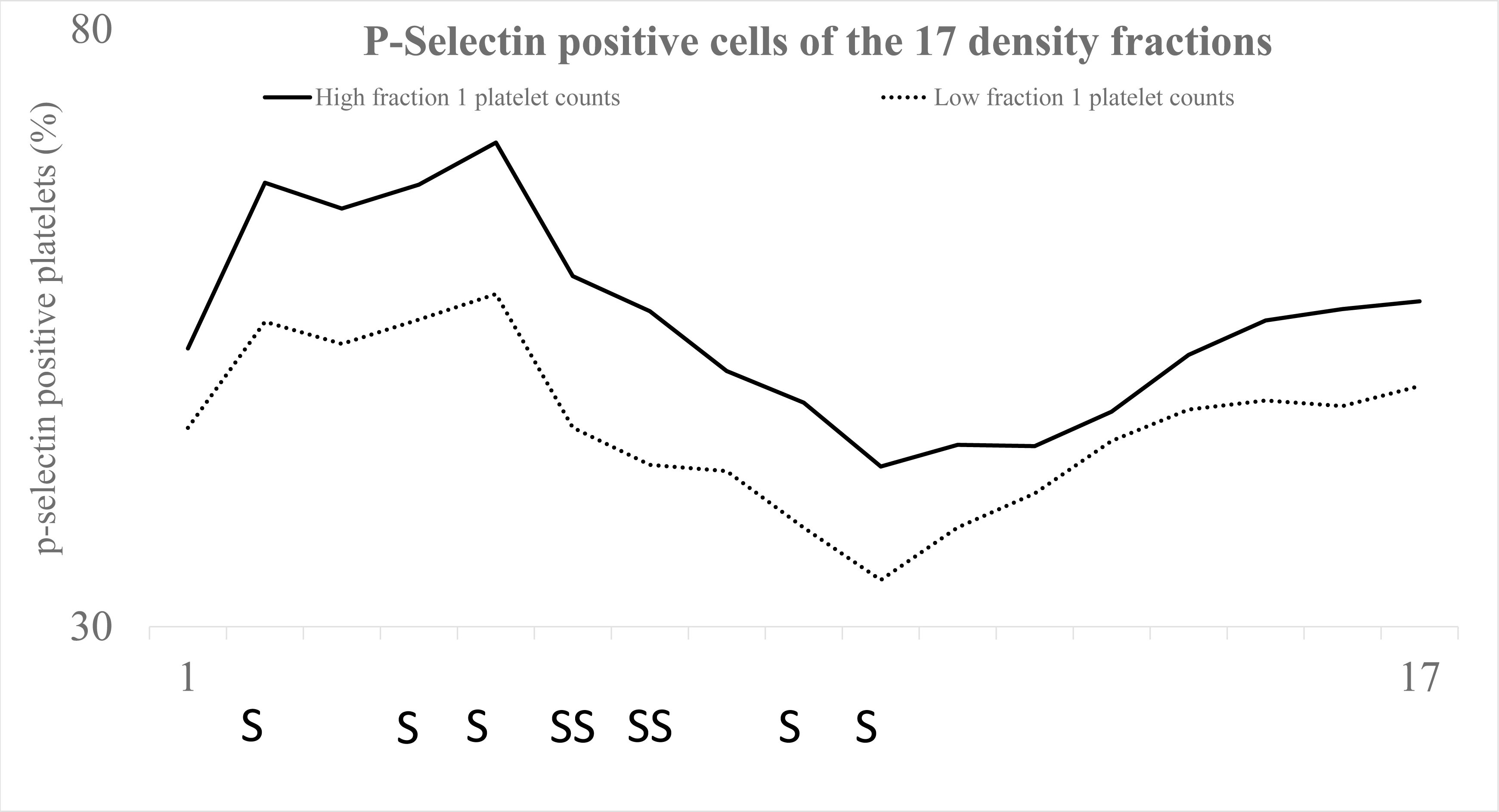
Displays, for the study groups (1.09^high^ and 1.09^low^), the percentage P-selectin positive platelets (vertical axis) of the 17 platelet fractions (horizontal axis). All platelet populations of 1.09^high^ individuals demonstrated elevated platelet P-selectin expression. Significances are denoted with the letters S (*p*≤0.05) and SS (*p*≤0.01). The differences proved to be significant for the fractions *nos*. 2, 4, 5 (*p*≤0.05), *nos.* 6, 7 (*p*≤.0.01) and *nos.* 9, 10 (*p*≤0.05).

**Figure 3.**
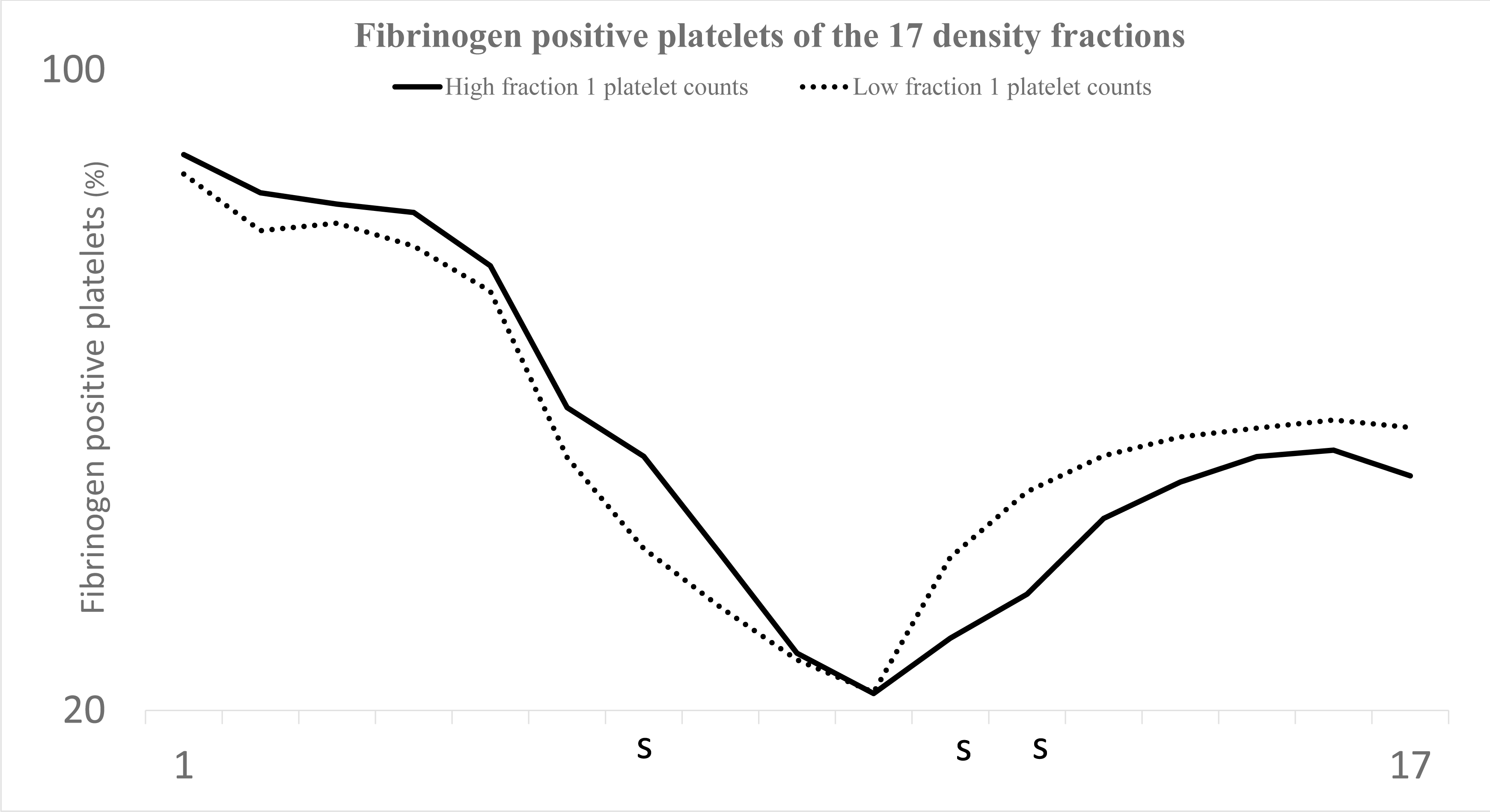
Results for the groups (1.09 ^high^ and 1.09 ^low^) with respect the percentage of fibrinogen-positive cells (vertical axis) of the density fractions (horizontal axis) (*n*=17) in the gradient. Significances are designated with the character S (*p*≤0.05). More dense platelets of 1.09^high^ individuals displayed more fibrinogen-positive cells and in fraction no. 7 the difference reached statistical significance (*p*≤.0.05). Lighter platelets of 1.09^high^ participants had less surface-bound fibrinogen and the differences were significant for the density fractions *nos.* 11, 12 (*p*≤0.05).

## Discussion

During the last few decades, interest in platelet heterogeneity has risen and considering their diversity it is anticipated that platelets execute diverse tasks in human physiology. This paper concludes that very high density (1.09 kg/L) platelets affect reactivity and activity of all platelets. Several lines of evidence support the notion most importantly that 1.09^high^ associates with membrane attached P-selectin after agonist provocation (reactivity) in whole blood. Furthermore, an inverse link was demonstrated to platelet surface fibrinogen *in vivo* (activity) (Table 2). Finally, 1.09^high^ individuals displayed many subpopulations with enhanced surface P-selectin (Figure 2) and they connect inversely with membrane attached fibrinogen of lighter cells (Figure 3).

According to previous work both high and low-density cells circulate more activated as judged by surface-bound fibrinogen [12], [21]. The findings are highly reproducible (Figure 4). Here a similar phenomenon was shown for platelet attached P-selectin. Very dense platelets (1.09 kg/L) however express less P-selectin (Figure 3) but more surface-bound fibrinogen. It is in keeping with the concept that platelets regulate α-granule release and GPIIb/IIIa receptor activation, independently [8]. Platelet functions affecting inflammation are favoured by surface P-selectin facilitating the recruitment of macrophages [7]. Thus, it is possible that just such very dense platelets are important in haemostasis thereby interfering less with inflammatory reactions.

Platelet heterogeneity has long been the subject of study and this laboratory has shown substantial alterations of the activation state of resting platelet subpopulations. 1.09^high^ individuals display an inverse relationship with whole blood reactivity and fibrinogen expression of lighter cells (Table 2) (Figure 3). It makes it feasible that the specific platelet phenotype has inhibitory effects upon aggregation. Very high-density cells reflect reactivity of all platelets (Table 2) but it is also evident that they move the entire density curve to the right (Figure 1) indicating less density of the whole platelet population. The findings disagree with the notion that density and reactivity are interrelated. Platelet activation, by necessity, involves granule release making cell density to decline. High density platelets demonstrate substantial activation as estimated from surface-attached fibrinogen (Figure 3) and such cells contain fewer α-granules [19]. Consequently, the subset with a high degree of certainty obtains its characteristics and abilities at thrombopoiesis because is unlikely that platelets acquire density when circulating. It is thus plausible that they are released very dense and designated to control reactivity of all platelets. It is possible, however, that alterations *in vivo* for instance through granule release are imposed upon an original hyper-density.

The current study investigates heterogeneity of unstimulated platelets and none of the populations demonstrate procoagulant features. Coated platelets are normally smaller displaying surface retention of α- granule proteins (i.e. P-selectin) but less GPIIb/IIIa activity [10]. Very dense (1.09 kg/L) platelets show increased GPIIb/IIIa receptor activation (Figure 2) but less P-selectin expression (Figure 3). Consequently, it is reasonable to assume that the current high-density population (1.09 kg/L) differs from procoagulant platelets and constitute an entity of its own.

## Conclusions

Putative roles of platelets are diverse and important interrogations in platelet diversity are relevant for human physiology. The subset of high-density (1.09 kg/L) procoagulant platelets has distinct properties, including the ability to influence reactivity of all platelets. Evidence supports the notion that platelets are inclined to certain tasks at their creation in the bone marrow.

## Acknowledgements

The Östergötland County Council, Sweden supported the research.

